# Single-cell RNA Sequencing Reveals Immunosuppressive Myeloid Cell Diversity and Restricted Cytotoxic Effector Cell Trafficking and Activation During Malignant Progression in Glioma

**DOI:** 10.1101/2021.09.24.461735

**Authors:** Sakthi Rajendran, Clayton Peterson, Alessandro Canella, Yang Hu, Amy Gross, Maren Cam, Akdes Serin-Harmanci, Rosario Distefano, Giovanni Nigita, Wesley Wang, Katherine E. Miller, Olivier Elemento, Ryan Roberts, Eric C. Holland, Ganesh Rao, Elaine R. Mardis, Prajwal Rajappa

## Abstract

Low grade gliomas **(LGG)** account for about two-thirds of all glioma diagnoses in adolescents and young adults (AYA) and malignant progression of these patients leads to dismal outcomes. Recent studies have shown the importance of the dynamic tumor microenvironment in high-grade gliomas **(HGG**), yet its role is still poorly understood in low-grade glioma malignant progression. Here, we investigated the heterogeneity of the immune microenvironment using a platelet-derived growth factor (**PDGF**)-driven **RCAS** (replication-competent ASLV long terminal repeat with a splice acceptor) glioma model that recapitulates the malignant progression of low to high-grade glioma in humans and also provides a model system to characterize immune cell trafficking and evolution. To illuminate changes in the immune cell landscape during tumor progression, we performed single-cell RNA sequencing on immune cells isolated from animals bearing no tumor **(NT)**, LGG and HGG, with a particular focus on the myeloid cell compartment, which is known to mediate glioma immunosuppression. LGGs demonstrated significantly increased infiltrating T cells, CD4 T cells, CD8 T cells, B cells, and natural killer cells in the tumor microenvironment, whereas HGGs significantly abrogated this infiltration. Our study identified two distinct macrophage clusters in the tumor microenvironment; one cluster appeared to be bone marrow-derived while another was defined by overexpression of Trem2, a marker of tumor associated macrophages. Our data demonstrates that these two distinct macrophage clusters show an immune-activated phenotype (*Stat1, Tnf, Cxcl9 and Cxcl10*) in LGG which evolves to an immunosuppressive state (*Lgals3, Apoc1 and Id2*) in HGG that restricts T cell recruitment and activation. We identified CD74 and macrophage migration inhibition factor (**MIF**) as potential targets for these distinct macrophage populations. Interestingly, these results were mirrored by our analysis of the TCGA dataset, which demonstrated a statistically significant association between CD74 overexpression and decreased overall survival in AYA patients with grade II gliomas. Targeting immunosuppressive myeloid cells and intra-tumoral macrophages within this therapeutic window may ameliorate mechanisms associated with immunosuppression before and during malignant progression.

## INTRODUCTION

Low-grade gliomas (**LGGs**) represent approximately one third of all CNS tumors [1]. In the adolescent and young adult (AYA) population, approximately two thirds of gliomas are low-grade [2] in this unique patient population [3]. Of note, two thirds of patients with LGG transform to high-grade gliomas in the IDH1 mutant setting (**HGG** - grade III or IV) [4], and the median overall survival after malignant progression is only 2.4 years [5]. To date, no approved therapies are available to prevent malignant progression in patients with low-grade glioma and radiologic imaging is the primary mode of gauging progression. [6]. The current standard of care consists of maximally safe resection followed by adjuvant therapy, including chemotherapy, radiation therapy, or combination chemoradiation therapy. However, these treatments are fraught with comorbidities and toxicities ranging from surgical complications, endocrine dysfunction, neurocognitive delay, and impaired neurologic function in the AYA population.

Although recent studies have shown LGG malignant progression is associated with overexpression of fibrinogen-like protein 2 in patients and murine models demonstrating increased regulatory T cells and M2 macrophage signatures, the molecular mechanisms of malignant progression require further elucidation [7]. Given the immunosuppressive tumor microenvironment (TME) within gliomas, current immunotherapy approaches with anti-PD1 immune checkpoint blockades have had limited success in improving patient survival in primary HGGs and recurrent glioblastoma multiforme (**GBM**) [8]. In 2020, one study described blocking the PD1 pathway, which promoted M1 macrophage polarization in murine glioma models independent of CD8 T cells [9]. Furthermore, tissue-specific deletion of PD1 on myeloid-derived cells enhanced antitumor immune responses in melanoma [10]. Moreover, the presence of myeloid-derived suppressor cells (**MDSC**) and tumor-associated macrophages negatively impact the efficacy of immune checkpoint blockade therapies by creating an immunosuppressive environment [11]. To that end, monocytic and granulocytic MDSCs have been shown to suppress T cell immune responses in murine GL261 glioma models [12].

We previously described the association of bone marrow-derived myeloid cell expansion, mobilization, and infiltration during LGG to HGG malignant progression using a platelet-derived growth factor (**PDGF**)-driven replication-competent ASLV long terminal repeat with a splice acceptor (**RCAS**) model [13, 14]. Various chemokines and cytokines secreted in the tumor microenvironment (**TME**) recruit myeloid-derived cells to the brain from peripheral circulation [14, 15]. Depleting macrophages by targeting CSF1R appeared to demonstrate preclinical efficacy in HGG models while modulating the Jak-Stat pathway and KDR/ID2 axis in myeloid cells appeared to impair LGG malignant progression in murine PDGF-RCAS LGG to HGG models [14, 16]. Despite these preliminary reports, there is still an unmet need to understand the functional diversity of myeloid cells within the TME. Identifying immunosuppressive myeloid cell vulnerabilities using single-cell RNA sequencing may provide therapeutic opportunities for patients with low-grade gliomas at risk of malignant progression. Here, we interrogated the heterogeneity within the myeloid compartment and its association with lymphoid cells in the TME during distinct points of LGG to HGG progression using single-cell RNA sequencing.

## MATERIALS AND METHODS

### Animals

The RCAS-Nestin tv-a (**Ntv-a**) model has been described before [17]. Ntv-a animals possess a transgenically expressed tv-a (receptor for RCAS) under the control of a Nestin promoter. Ntv-a animals were crossed with Ink4a-Arf-/- LPTEN animals. The heterozygous pups were injected with 100,000 cells of DF-1 cells transfected with RCAS-Cre and RCAS-PDGFB in the brain parenchyma using a 10 μl gastight Hamilton Syringe. The Institutional Animal Care and Use Committee (IACUC) of Nationwide Children’s Hospital (Protocol number: AR19-00146) approved all animal experiments.

### Tissue processing and H&E staining

After euthanizing the animals in a CO_2_ chamber, the brains were extracted and fixed in 10% neutral buffered formalin, followed by embedding in paraffin and sectioning at 4 µm. Slides were stained with hematoxylin and eosin. Images were acquired using Aperio Scanscope and processed using Imagescope.

### Cell culture

DF-1 cells were cultured in DMEM with 10% fetal bovine serum (**FBS**) and antibiotics (Penicillin and Streptomycin) at 37°C. DF-1 cells were transfected with vectors RCAS-Cre and RCAS-PDGF using Fugene.

### Single-cell preparation

Tumor or similar-sized normal tissue was excised from the brain’s right hemisphere, where the cells had been injected. The tissue was minced, triturated, and digested with 2 mg of Papain (Brainbits, Cat. # PAP/HE) for 20 minutes at 37°C. After digested tissue was triturated and passed through a 70 µm cell strainer and CD45+ cells were isolated with a Miltenyi positive selection kit (Cat. # 130-052-301), according to the manufacturer’s instructions.

### Single-cell RNA sequencing

Single cell suspensions were run on a Chromium controller (10x Genomics) using the chromium Next GEM Single Cell 3’ Reagent kit v3.1, targeting a capture of 5000 cells. Sequencing results were mapped to the mm10 mouse genome reference build and quantified using the CellRanger v3.0.2 (10xGenomics) [18, 19] (https://support.10xgenomics.com/single-cell-geneexpression/software/pipelines/latest/installation). Cells (n= 14,125) were identified as HGG (n=4468), LGG (n=5844), and no tumor (**NT**) (n=3813). Seurat v.4 and ShinyCell packages were utilized to evaluate gene expression analysis in R [19-21]. We excluded cells with fewer than 200 transcripts or more than 10% mitochondrial genes to filter out low-quality data. With these filters, 12,668 cells remained for further analysis (HGG=4100, LGG=5024, and NT=3544 cells). Gene expression measurements were LogNormalized, and the data were scaled, regressing out the transcript counts and mitochondrial gene percentage. The “mean.var.plot” method revealed variable features. After integrating the data, principal component analysis using the first 38 principal components for further analysis reduced the dimensionality. Unsupervised clustering was performed with the resolution set to 0.8. Two-dimensional t-SNE and Uniform Manifold Approximation and Projection (**UMAP**) approaches visualized the clustering results [22, 23]. Canonical marker expression identified and further analyzed immune cluster types. Cell counts determined the cells in each cluster at each tumor time. The Wilcoxon rank-sum test (min.pct = 0.1) identified differentially upregulated genes between groups and functionally analyzed clusters. Differentially upregulated and downregulated genes were also identified with the Wilcoxon rank-sum test (min.pct = 0.1, only.pos = FALSE) and used for IPA analysis (Qiagen).

### Random forest modeling

Random forest modeling was performed utilizing the random Forest package in R [24]. Normalized expression counts were extracted from the scRNA-seq Seurat object, along with cluster labels and cell barcodes. Information was merged into a novel data frame in R composed of a cell’s cluster, normalized gene expression values, and the cancer progression state the cell was extracted from [Normal, LGG, HGG]. To reduce computational complexity, evaluated genes for each cell cluster were filtered over genes identified as differentially expressed from our above analysis and split into individual data frames composed of cells from each cluster. Data was then passed through Boruta to further reduce features (genes) for modeling (**Supplementary Figure 5**) [25]. Utilizing the reduced feature list, random forest models were generated to predict a cell’s cancer progression state origin utilizing gene expression profiles. Data was split 60% for model training and the remaining 40% for validation testing. Validation performance was evaluated using caret and pROC to calculate confusion matrix scores and receiver operating characteristic-area under the curve (AUC) [26, 27]. Performance metrics and ranked importance were visualized using ggplot2 [28].

### Ingenuity Pathway Analysis (**IPA**) analysis

IPA (QIAGEN Inc., https://www.qiagenbioinformatics.com/products/ingenuity-pathway-analysis) was used to generate network analysis, and upstream predicted targets [29].

### Flow cytometry

Tumor tissue excised from the brain was mechanically dissociated using a 70 um cell strainer in phosphate-buffered saline (**PBS**) with 10% FBS and 0.1 mg/mL DNAse I (Stem Cell Technologies). Cells were then resuspended in 25% isotonic Percoll solution and centrifuged for 20 minutes at 18°C with acceleration and deceleration set to minimum. After removing myelin debris and Percoll, the cell pellet was washed with 10 mL of PBS with 10% FBS. Cells were resuspended in 1x PBS and stained with the Invitrogen™ LIVE/DEAD™ Fixable Blue Dead Cell Stain Kit. Cells were then washed and blocked with 1:100 of FcR blocking solution (BD Biosciences) for 10 minutes on ice. Cells were stained with CD11b-BV421, CD3-PE-Cy7, CD8-APC-Cy7, F4/80-APC, CD45-PE, and Gr1-FITC, then washed and fixed with 1% PFA for 5 minutes on ice. Flow cytometry data were acquired using BD Fortessa X-20 and analyzed using FlowJo-v10.

### Quantitative polymerase chain reaction (**qPCR**)

Myeloid and Lymphoid tumor-infiltrating populations were isolated as previously described (flow cytometry method). RNEASY Micro Kit (74004, Qiagen) was used to extract total RNA, and the High Capacity Reverse Transcription Kit (4387406, Applied Biosystems) was used to synthesize cDNA. qPCR product was amplified using SYBR Green master mix (A25742, Applied Biosystems) and StepOnePlus (Applied Biosystems). The primers (IDT) used included the following: CD74 (Fwd: CTGATGCGTCCAATGTCCAT; Rev: CAGACCTCGTGAGCAGATG), ID2 (Fwd: CACTATCGTCAGCCTGCATC; Rev: TCATTCGACATAAGCTCAGAAGG), GAPDH (Fwd: AATGGTGAAGGTCGGTGTG; Rev:GTGGAGTCATACTGGAACATGTAG), TBP (Fwd: TGTATCTACCGTGAATCTTGGC; Rev: CCAGAACTGAAAATCAACGCAG).

### Tissue lysate preparation

After excising, tumor or normal tissue was homogenized in a pyrex tissue grinder with 1 mL of tissue extraction buffer. The Pierce BCA protein assay kit was used to quantify total protein. Samples were reconstituted to a final protein concentration of 5 µg/mL and stored at −80°C. Chemokines were quantified using the Luminex kit by Eve Technologies.

### Survival analysis (TCGA dataset)

RNA-seq expression levels were extracted from The Cancer Genome Atlas Low Grade Glioma project data (TCGA-LGG - https://www.cancer.gov/tcga) as fragments per kilobase of transcript per million mapped reads (FPKM) using the Genomic Data Commons Portal [30, 31]. As defined by TCGA, “lower grade glioma” samples included both grade 2 and grade 3 gliomas (astrocytoma, oligodendroglioma, and mixed gliomas). Samples were filtered for cases less than 40 years of age and stratified by grade. Genes with FPKM geometric means greater than 1 across patients were retained and then raw counts were normalized using Trimmed Mean of M-values. Values were calculated for genes of interest and patients from the upper (n=34) and lower quartiles (n=34) of expression were plotted for survival and compared using a Log-rank test P-value. Accompanying clinical data and molecular markers describing the assayed patient populations were extracted using TCGAbiolinks and summarized in **Supplementary Table 1** [32-35].

### Survival analysis (Glioma dataset)

Paired-end RNA sequencing was performed on total RNA extracted from formalin-fixed, paraffin-embedded (FFPE) tissues from secondary GBM (Grade III (n=44), Grade IV (n=23)) and LGG (grade II, n=42) using MasterPure kit (Epicentre). Raw reads from the RNA-seq data were processed using an in-house pipeline that uses STAR for read alignment; FastQC and RSeQC (for read and alignment quality assessment) and FeatureCount for expression count. Samples with degraded RNA or unique mapping rates lower than 30% were excluded. The reads were aligned to GRCh38 Human reference genome and mapped to the human transcriptome according to UCSC gene annotations. We then normalized the RNA-seq read counts for genes and applied a variance stabilizing transformation. We transformed the expression data to z-scores and set a Z-scale cutoff for high and low expression. Expression values higher than 0 are set to “high” and lower than 0 are set to “low” and compared these against “overall survival” in a Cox Proportional Hazards (Cox) survival model. We included sample grade information as a covariate in the model. We used survminer R package for the survival analysis.

## RESULTS

### Single-cell RNA sequencing identifies quantifiable differences in immune cell phenotypes during glioma progression

Immature myeloid-derived cells may restrict T cell infiltration and activation, leading to immunosuppression in gliomas [36]. We are specifically interested in studying the heterogeneity of myeloid cell populations and the functional consequence of this diversity in the TME during LGG malignant progression. To that end, we have used a murine PDGF-B driven RCAS glioma model [16, 17] that develops LGGs starting around 3 to 4 weeks and progresses to high-grade glioma at about 6 to 8 weeks (**Figure 1A**). Here, we performed flow cytometry for CD11b and Gr1 (Ly6c/Ly6g), a murine marker of myeloid-derived suppressive cells in the bone marrow of tumor-bearing animals at specified times. The proportion of CD11b+Gr1+ cells significantly expanded in the bone marrow of animals between 3-4 weeks and in HGG (39.4 ± 2.7%) when compared to LGG (18.7 ± 2.1%) (**Supplementary Figure 1**). Increased CD11b+Gr1+ cells correlated with tumor progression and aligns with other reports [14, 37], however our observation details the initial timing for bone marrow expansion of MDSC-like cells.

**Figure 1.**
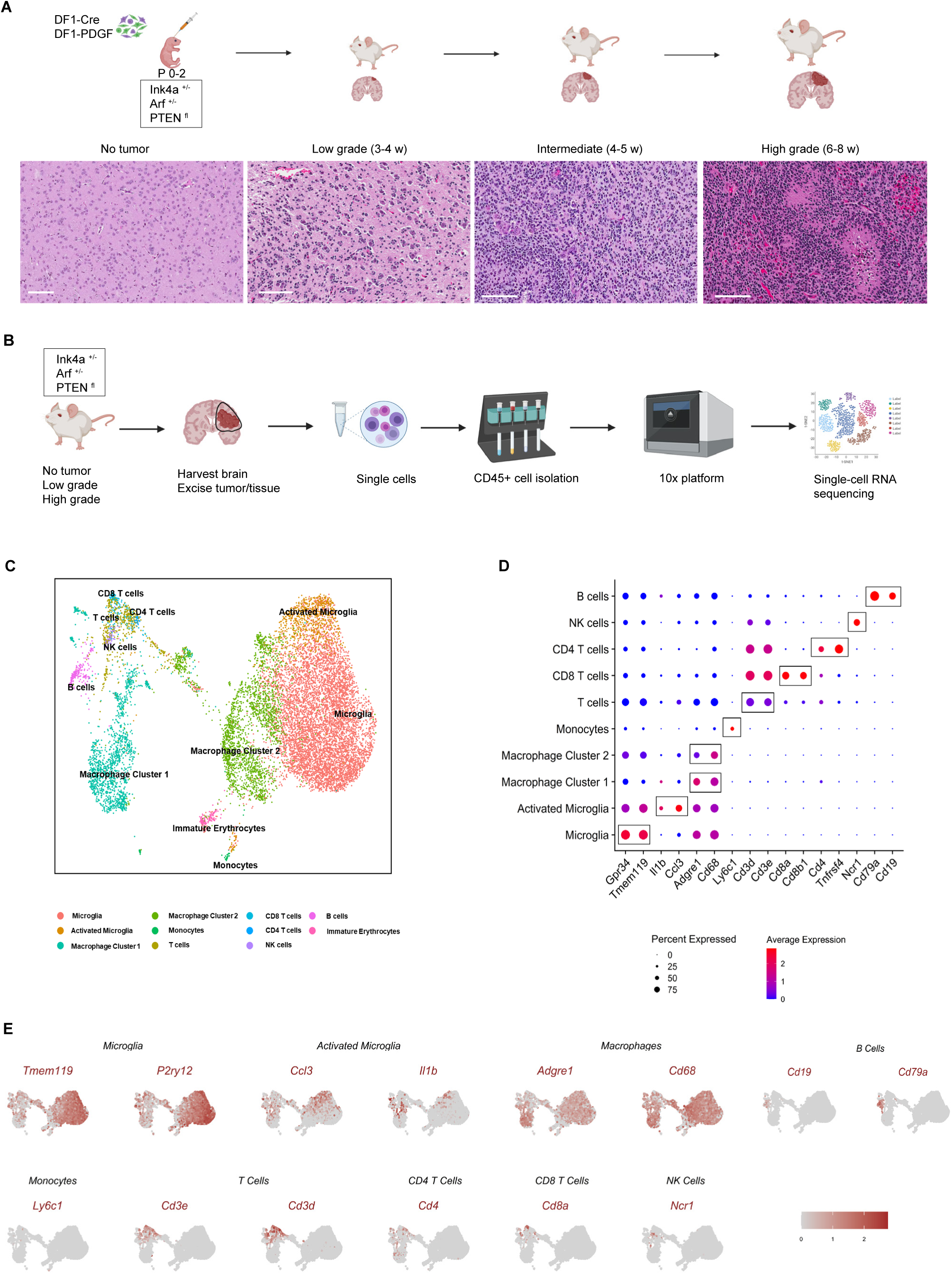
Single-cell RNA sequencing of tumor-infiltrating leukocytes during malignant progression. (**A**) A schematic representing the platelet-derived growth factor (**PDGF**)-driven replication-competent ASLV long terminal repeat with a splice acceptor (**RCAS**) glioma progression from low-grade (3-4 w) to high-grade (6-8 w) with representative H&E stained images. Scale bar – 100 µm. (**B**) Graphic illustrating single-cell RNA sequencing from CD45+ cells (lymphocyte common antigen) isolated from brains of animals with no tumor (**NT**) (n=2), low-grade glioma (**LGG**) (n=3), and high-grade glioma (**HGG**) (n=3). (**C**) Uniform Manifold Approximation and Projection (**UMAP**) for dimensionality reduction representing the cell clusters of the integrated dataset of all samples obtained using unsupervised clustering in Seurat. (**D**) Dot plots demonstrating canonical gene marker expression for each cell cluster. (**E**) Feature plots showing canonical marker expression for each cluster in the UMAP.

We expanded upon these results by performing single-cell RNA sequencing on isolated CD45+ cells from pooled tumor or normal tissues from animals with no tumor (NT, n=2), LGG (n=3), and HGG (n=3) using the 10x Genomics platform (**Figure 1B**). We analyzed 12,668 cells from the three samples after quality control, then integrated data from all samples and projected the data using UMAP (**Figure 1C**). Unsupervised cell clustering was performed using Seurat V4. We identified a total of 11 distinct clusters of known immune cells comprising disparate populations of microglia, macrophages, monocytes, T cells, B cells, natural killer cells and immature erythrocytes (**Figure 1C**). Cell cluster identities were verified by expression of canonical markers for each cell type. These cell cluster datasets are visualized using dot plots (**Figure 1D**) and feature plots (**Figure 1E)**.

To identify immune cell landscape dynamics during glioma progression, we calculated the ratio of each cell cluster per total cells analyzed in each sample, then evaluated the myeloid and lymphoid cell distribution and heterogeneity across NT, LGG, and HGG **(Figure 2A)**. We observed a significantly increased proportion of the total macrophages and a significantly reduced T cell proportion in HGG compared to LGG (**Figure 2B**). In order to validate these results, we performed flow cytometry on dissociated brain tumor tissues of LGG and HGG. We quantified the percentages of CD45 high and CD11b+ cells, conventionally identified as bone marrow-derived myeloid cells **(BMDM)** [38, 39]. Using flow cytometry, we detected a 4-fold *in vivo* increase in the percentage of BMDM in HGG (11.5 ± 4.5%) compared to LGG (2.7 ± 1.5%) (**Figure 2C and 2D)**. These CD45high and CD11b+ cells also expressed F4/80, a macrophage marker, in 63.5 ± 7% of LGG and in 87.3 ± 5.1% of HGG (**Figure 2C and 2D)**. In CD45 high and CD11b+ cells, the difference in CD11b+Gr1+ cells (myeloid-derived suppressive cells) between LGG (5.5 ± 3.2%) and HGG (10.9 ± 4.3%) animals was not statistically significant (**Figure 2D)**. In addition, we validated the presence of T cells in animal brains with low-grade and high-grade glioma by flow cytometry **(Figure 2E)**. We found significantly increased T cells in the lymphocyte population (CD45^hi^CD11b+) in LGG (78.5 ± 1.4%) than in HGG (43.4 ± 11.3%). Similarly, LGG had significantly higher proportion of CD8+ T cells (33 ± 6.2%) than HGG (8.6 ± 10.9%) **(Figure 2F)**. These results support our single-cell RNA sequencing data which revealed significantly reduced T cell infiltration in HGG.

**Figure 2.**
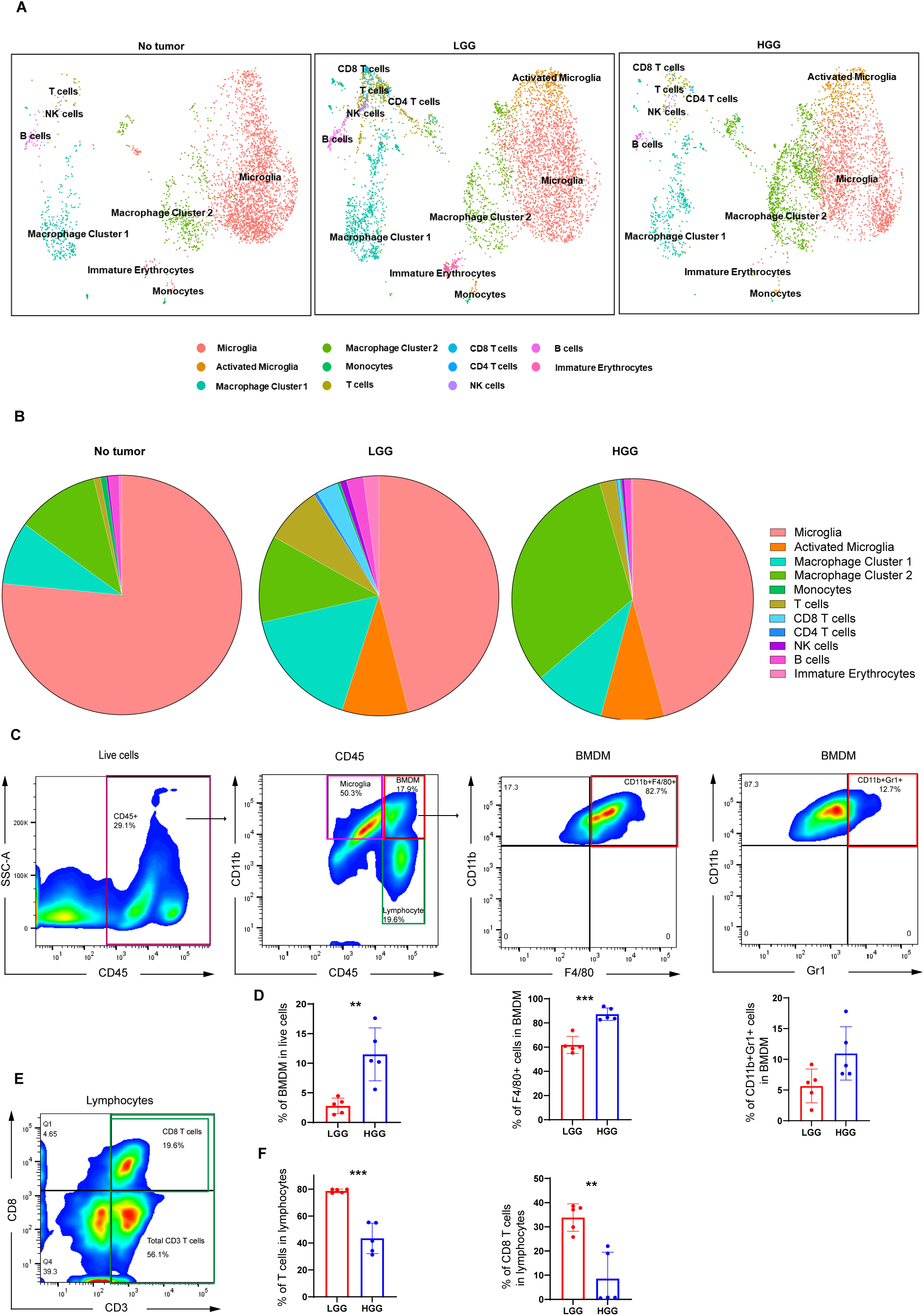
Single-cell RNA sequencing reveals differences in immune cell infiltration during glioma progression. (**A**) Uniform Manifold Approximation and Projection (**UMAP**) representing immune cell clusters in samples with no tumor, low and high-grade gliomas. (**B**) Pie charts illustrating the proportion of immune cell clusters identified in each sample. (**C**) Flow cytometry gating strategy for BMDM and lymphocytes in high-grade glioma (**HGG)** and dot plots representing F4/80 or Gr1 staining in BMDM **(D)** Quantification of CD11b+F4/80+ cells and CD11b+Gr1+ in BMDM in animals bearing low-grade glioma (**LGG**) (n=5) and HGG (n=5). **(E)** Representative dot plot showing flow cytometry staining of CD3 and CD8 in lymphocyte population in an animal with HGG. **(F)** Quantification of Total CD3 T cells and CD8 T cells within lymphocytes of animals bearing LGG (n=5) and HGG (n=5). A two-tailed unpaired student’s t-test was used to calculate statistical significance. *P<0.05, **P<0.01, ***P<0.005.

### Diversity in microglial populations during tumor progression

The predominant immune cell type observed in the normal brain was microglia, with 76.6% of all immune cells clustered as microglia. In contrast, only 45.8% and 46.1% of immune cells in LGG and HGG were microglia, respectively (**Figure 2A, 2B, Supplementary Figure 2A, 2B)**. Strikingly, the activated microglial population was absent in the normal brain and was observed in only LGG and HGG (9% and 8.5% of all immune infiltrates, respectively), **Figure 2A** and **2B**. Activated microglia also significantly expressed more chemokines (e.g., *Ccl4, Ccl3, and IL-1b)* and other signatures of transcriptional activation (e.g., *Jun, Junb, Junc, Fos, Fosb, and Socs3*) than microglia (**Supplementary Figure 2C**), as also identified by this report [40].

### Macrophages adopt immunosuppressive transcriptional signatures during malignant progression

Single-cell RNA sequencing identified two distinct macrophage clusters. We observed an increase of macrophage cluster 1 in LGG (16.5%) relative to normal brain (8.5%) and to HGG (9.5%), **Figure 2A, 2B**. Macrophage cluster 1 also expressed unique signatures of bone marrow-derived macrophages (**BMDM**) (e.g., *TgfbI, Cd14, and Itga4*) (**Figure 3A**), as described by these reports [41, 42]. By comparison, Macrophage cluster 2 expressed signatures of Trem2 high macrophages (e.g., *Cd9, Cd81, Ctsb, Ctsd, and Ctsd*) (**Figure 3A**). This cluster also demonstrated significantly downregulated signatures of BMDM-associated genes while overexpressing *Trem2* (**Figure 3A)**. In addition, Trem2 is a marker of tumor-associated macrophages [43], and increased Trem2 hi macrophages have been reported in non-responders to immune checkpoint inhibitors in patients with melanoma [44].

**Figure 3.**
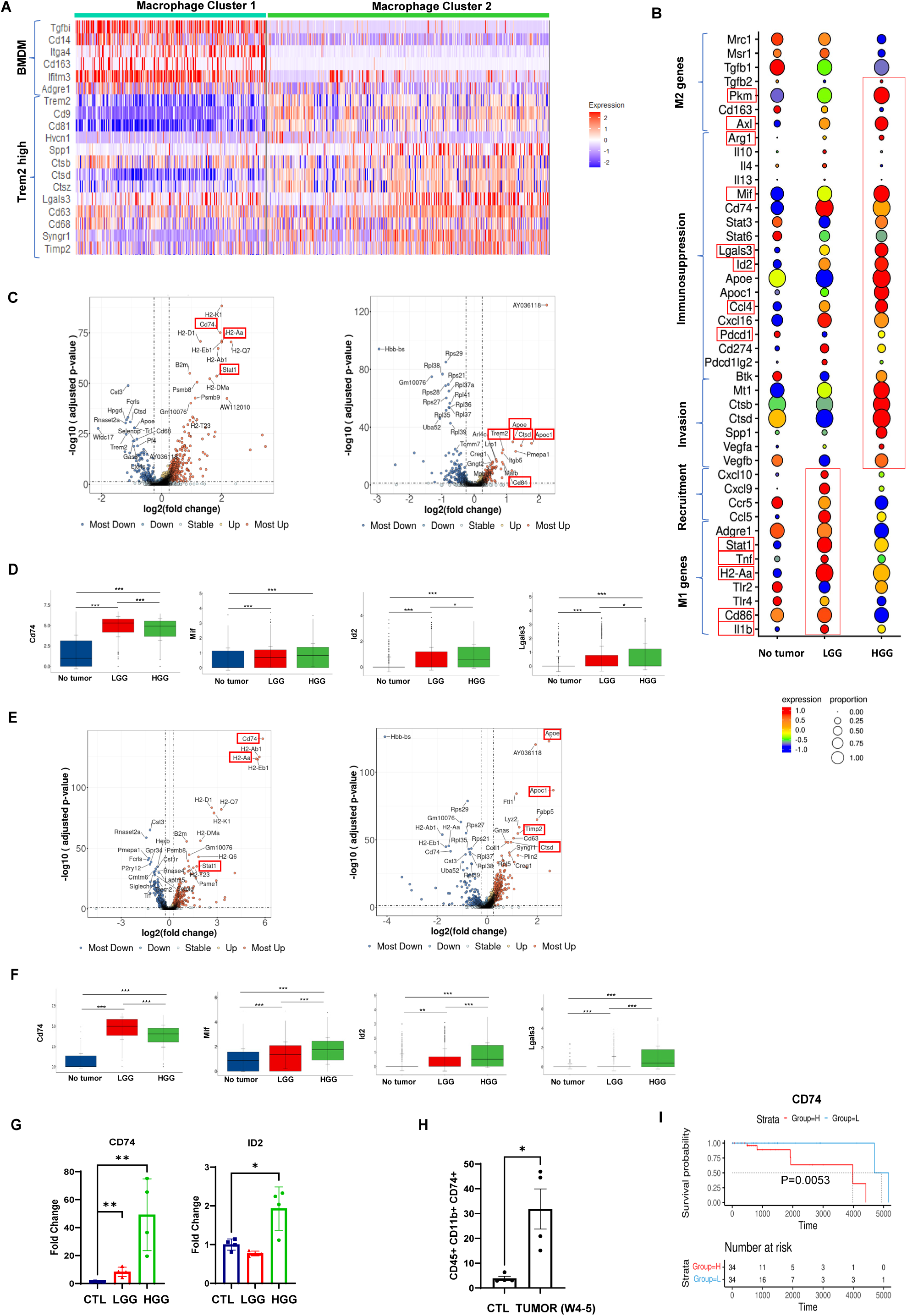
Heterogeneity in macrophage clusters during glioma progression. (**A**) Heatmap of signature genes of bone marrow-derived macrophages (**BMDM**) and Trem2 high macrophages in macrophage clusters 1 and 2. (**B**) Differential expression analysis of genes associated with M2 polarization, immunosuppression, invasion, recruitment, and M1 polarization in combined macrophage cluster. (**C**) Volcano plots showing top 15 differentially expressed genes in macrophage cluster 1 in no tumor (**NT**) to low-grade glioma (**LGG**) and LGG to high-grade glioma (**HGG**). **(D)** Box plots representing the expression of key immunosuppressive molecules (*Cd74, Mif, Id2* and *Lgals3*). **(E)** Volcano plots representing the top 15 differentially expressed genes in macrophage cluster 2 in NT to LGG and LGG to HGG, respectively. **(F)** Box plots representing the expression of key immunosuppressive molecules (*Cd74, Mif, Id2* and *Lgals3*) in macrophage cluster 2. **(G)** qPCR analysis for *Cd74* and *Id2* in animals with NT (n=4), LGG (n=4) and HGG (n=4). **(H)** Flow cytometry analysis of CD74 staining in tumor infiltrating bone-derived myeloid cells of animals with NT (n=4) and tumor (n=4). **(G)** Overall survival of grade II glioma patients from the TCGA dataset expressing low levels (blue) and high levels (red) of *Cd74*. A two-tailed unpaired student’s t-test was used to calculate statistical significance. *P<0.05, **P<0.01, ***P<0.005.

Integrated data revealed both macrophage clusters overexpressed significantly increased immunosuppressive factors (e.g. *Arg1, Mif, Lgals3, Pkm, Axl, Btk, Id2, Ccl4, Pdcd1*) in HGG compared to LGG (**Figure 3B**). In contrast, LGG had significantly increased levels of recruitment factors and M1 genes (*Stat1, Tnf, HLAII (H2-Aa), Cd86, and IL-1b*) (**Figure 3B**). This trend also manifests in macrophage clusters 1 and 2, respectively (**Supplementary Figure 2D, 2E)**. Overall, our data suggest that both macrophage clusters develop immunosuppressive signatures as LGGs progress to HGG.

### Heterogeneity and distinct macrophage cluster function during malignant progression

Macrophage cluster 1 significantly expressed more CD74, MHCII (H2-Aa) genes, and *Stat1* in LGG than NT (**Figure 3C**). Macrophage cluster 1 from HGG had significantly increased expression of *ApoE, ApoC1, Trem2, Cd81, and Ctsd* than LGG (**Figure 3C**). The top 20 differentially expressed genes in this macrophage cluster included molecules involved in immunosuppression (e.g., *Apoc1, Apoc4, ApoE, Spp1*), and matrix remodeling factors such as *Ctsd* (Cathepsin) and *Itgb5* (Integrin) in HGG (**Supplementary Figure 3A**). ApoE expression in macrophages downregulates pro-inflammatory M1 factors and induces an immunosuppressive M2 phenotype expressing *Arg1* [45]. In contrast, macrophage cluster 1 in LGG expressed T cell-recruiting chemokines (*Cxcl9, Ccl5, Ccl8*) and immune-activating signatures [e.g., *Stat1*, MHC-II molecules (*H2-Aa, H2-Q7*), and MHC-I gene (*H2-K1*)] (**Supplementary Figure 3A**). In addition, machine learning-based modeling using a random forest approach **(Supplementary Figure 5)**, demonstrated statistically significant predictive capability to discern the evolution of select cell clusters during tumor progression. Of note, this classifier algorithm demonstrated high predictive value for macrophage cluster 1 and its immunosuppressive evolution during tumor progression based on its transcriptional profile [Accuracy: 89%; AUC: 0.90] (**Supplementary Figure 6D)**. When we evaluated the predominant features (genes) that most strongly drove the model’s predictive success, we identified several immunosuppressive (Cd74, ApoE) and immune-activating or -recruiting (MHC genes, *Stat1, Cxcl9, Ccl8*) signatures (**Supplementary Figure 6A**). LGG and HGG samples upregulated more *Cd74* and its binding partner *Mif*, as well as *Id2 and Lgals3* (**Figure 3D**), compared to NT. Further, IPA analysis identified significantly increased expression of Stat1 and interferon-related genes in LGG compared to NT **(Supplementary Figure 3B and 3C)**. Macrophage cluster 1 in HGG downregulated more Ifng and Stat1 pathways than LGG (**Supplementary Figure 3D and 3E)**. This analysis also predicted the top upstream targets for activation in macrophage cluster 1: PPARG, PTGER4, and IL10RA, and downregulated pathways included TLR4, IFNAR1, TNF, IFNB1, STAT1, AND IFNG (**Supplementary Figure 3D and 3E)**.

Macrophage cluster 2 also significantly overexpressed Cd74 and MHCII (H2-Aa) genes in LGG relative to NT (**Figure 3E**). In HGGs, *ApoE, ApoC1, Timp2, Cd63*, and *Ctsd* were upregulated when compared to LGG (**Figure 3E)**. Among the top 20 differentially expressed genes, we identified immunosuppressive factors [e.g., *Lgals3* (Galectin-3), *Spp1, Pkm, ApoE, ApoC1*, and *Gpnmb*] in HGG (**Supplementary Figure 4A**). In contrast, LGG samples expressed more chemokines (*Cxcl10, Cxcl9, Ccl5*, and *Ccl8*) and HLA II genes (*H2-Aa, H2-Ab1, H2-Q6, and H2-Oa*) than NT and HGG (**Supplementary Figure 4A**). Furthermore, *in-silico* modeling using a random forest classifier algorithm showed high accuracy in predicting which state of progression a cell from macrophage cluster 2 originated from based on transcriptional profile alone [Accuracy: 89%; AUC: 0.89] (**Supplementary Figure 6E**). Furthermore, the key features used to accurately predict progression state included cluster defining genes such as *Timp2* and *Cd63*, but immunosuppressive markers *ApoE* and *ApoC1* were among the top 3 genes which drove decision making of the model (**Supplementary Figure 6B**). Similar to cluster 1, macrophage cluster 2 expressed significantly higher levels of *Cd74, Mif, Id2* and *Lgals3* in tumor samples than NT, P<0.05 (**Figure 3F**). IPA analysis identified macrophage cluster 2 in LGG expressed increased TNF, IFN-g, TLR4 and STAT1 compared to NT (**Supplementary Figures 4B and 4C**). By contrast, HGG downregulated IFNg and upregulated CSF1 and CD300LF relative to LGG (**Supplementary Figure 4D**). Both macrophage clusters showed upregulated expression of M1-related genes STAT1, TNF, TLR4, and IFN-g in LGG, but relative downregulation of TGFB, PTGER4, and IL10RA compared to NT **(Supplementary Figures 3B, 4B)**.

We identified *Spp1* among the top 20 upregulated genes in both the macrophage clusters in HGG (**Supplementary Figures 3A and 4A**). Likewise, *Gpnmb* was among the top upregulated genes in macrophage cluster 2 in HGG (**Supplementary Figure 4A**). Both *Spp1* and *Gpnmb* are markers of glioma-associated macrophages in both murine glioma models and patients with GBM [43]. Furthermore, patients with GBMs expressing higher levels of *Spp1* and *Gpnmb* had decreased overall survival compared to GBM patients expressing low levels of *Spp1* and *Gpnmb* [43]. In addition, inhibiting Spp1 expression impaired progression of tumors overexpressing MMP2 and Vimentin in a murine xenograft glioma model [46].

Given both macrophage clusters appear to demonstrate significant immunosuppressive properties, we identified common LGG markers that warrant further functional investigation. We observed LGG and HGG samples but not NT controls overexpressed Cd74 and Mif in both macrophage clusters, (**Figures 3D and 3F)**. We then validated the expression of *Cd74* and *Id2* by qPCR in cells isolated from the TME. We found significantly higher *Cd74* and *Id2* expression in HGG compared to LGG (**Figure 3G**). Flow cytometry staining also indicated HGGs overexpressed CD74 in CD45hi/CD11b+ cells (31.9 ± 16.2%) when compared to LGG (3.9 ± 1.7%) (**Figure 3H**). Interestingly, overexpression of *Cd74, Mif, and Spp1* within macrophages of HGG is associated with reduced overall survival in grade II glioma patients in TCGA datasets **(Figure 3I, Supplementary Figure 7A and 7B)**. Likewise, overexpression of *Lgals3* and *Ptger4* negatively correlated with overall survival in TCGA patients less than 40 years of age with low- and high-grade glioma **(Supplementary Figure 7C and 7D)**.

### Restriction of T cell trafficking and activation in the HGG TME

We observed a significant infiltration of T cells in LGG compared to HGG samples (**Figure 2B and Supplementary Figure 2A**). Total CD3+ T cells accounted for 7.8% of immune infiltrates in LGG, whereas normal brain and HGG had less than 1% of CD3+ T cells in each sample (**Figure 2B and Supplementary Figure 2A**). To that end, differential expression analysis identified an upregulation of chemoattractants such *as Ccl8, Cxcr6, Ifng*, and *Ccl5* in infiltrating T cells in LGG. In contrast, immune infiltrating T cells in HGG expressed more molecules associated with lipid metabolism (e.g., Apoe, Apoc1, Hmox, and Fabp5) (**Figure 4A and Supplementary Figure 8**).

**Figure 4.**
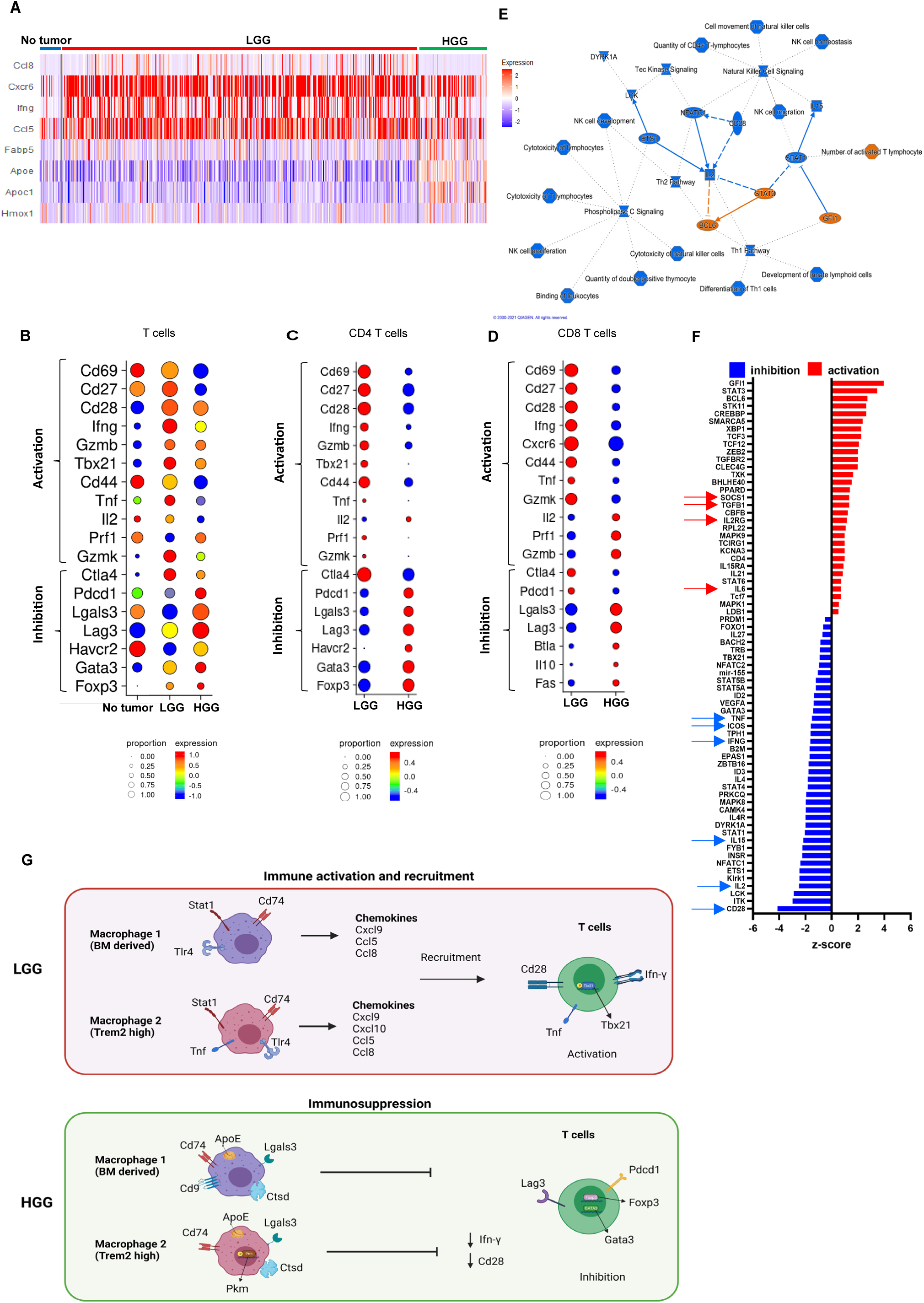
Impaired T cell activation in animals bearing HGG. (**A**) Heat maps showing key immune activation (*Ccl8, Cxcr6, Ifng*, and *Ccl5*) and suppression genes (*Apoe, Apoc1, and Hmox1*) in total T cells from samples with no tumor (**NT**), low-grade glioma (**LGG**), and high-grade glioma (**HGG**). Differential expression analysis showing genes related to T cell activation and inhibition in total CD3+ T cells (**B**), CD4 T cells (**C**), and CD8 T cells (**D**). **(E)** Network of pathways inhibited (blue) and activated (orange) in total T cells during progression from LGG to HGG. (**F**) Top upstream predicted targets in T cells during progression from LGG to HGG. (**G)** Schematic representation of crosstalk between macrophages and T cells in glioma tumor microenvironment indicating immune activation and suppression during LGG and HGG, respectively.

Then we investigated the differences in total CD3+ T cells between LGG and HGG. Total T cells in HGG expressed significantly fewer T cell activation markers (e.g., *Cd69, Ifn-g, Cd27, Cd44, Tnf*, and *Il-2*) than in LGG (**Figure 4B**). Tumor cell lysates in LGG also secreted more IL-2, Eotaxin, IL-1b, MCP-5, CXCL9 or MIG than NT and HGGs (**Supplementary Figure 9A)**. These cells also expressed increased immune checkpoint molecules PD1 (*Pdcd1*), *Lag3*, and increased Th2 transcription factor, *Gata3*, and T reg transcription factor, *Foxp3* (**Figure 4B**). Despite not detecting any CD4 and CD8 T cells in the NT sample, we found substantially more CD8+ T cells (2.9%) in LGG than in HGG (0.3%) (**Supplementary Figure 2A**). We found similar proportions of CD4+ T cells in LGG (0.5%) and HGG (0.2%) (**Supplementary Figure 2A**). CD4+ T cells expressed significantly higher levels of activation markers (e.g., *Cd69, Cd27, Cd28, Ifn-g, Gzmb*, Th1 transcription factor *Tbx21, and Tnf*) in LGG, and all these molecules were expressed significantly less in HGG (P<0.05). Likewise, T cell inhibitory markers *Pdcd1, Lgals3, Lag3, Havcr2 (Tim3)*, Th2 transcription factor (*Gata3*), and T reg marker (*Foxp3*) expression increased in T cells after malignant progression (**Figure 4C**). In CD8+ T cells, we observed *Cxcr6, Cd44, Cd69, Cd28, Cd27* in LGG and increased *Lgals3, Lag3, Btla4, Il10, and Fas* (**Figure 4D**). LGG had expressed significantly more *Cxcl9, Cxcl10, Ccl5, Ifng, and Cxcr6* than NT and HGG (P<0.05) (**Supplementary Figure 9B**). CXCR6 is a critical factor for homing and recruitment of CD8+ T cells [47]. IPA analysis also identified suppression of IL-2, Th1 polarization, and T cell and natural killer cell cytotoxicity (**Figure 4E**). Costimulatory molecules CD28, ICOS, IL-2, IL-15, IFNg, and Tnf were among the top downregulated predicted pathways. SOCS1, TGFB1, IL2R, and IL-6 were among the top upregulated pathways in HGG (**Figure 4F**). Interestingly, while the predictive capability of discerning the progression state of a T cell was high with a random forest model [Accuracy: 90%; AUC: 0.81] (**Supplementary Figure 6F**), the driving features of the model were not largely involved by many of the above-mentioned markers (**Supplementary Figure 6**). These results support that the immunosuppressive macrophage clusters within the TME may have restricted T cell function and infiltration as the tumor progressed to high-grade glioma **(Figure 4G)**.

## DISCUSSION

We investigated the immune cell landscape and heterogeneity of the myeloid compartment and its association with cytotoxic effector cell trafficking and activation during malignant progression of glioma using the PDGF-driven RCAS LGG to HGG model in adolescent and young adult mice. Using scRNAseq, our data revealed distinct differences in macrophage immune activation status during tumor progression, facilitating the identification of potential therapeutic targets specific to macrophages. We identified two distinct macrophage clusters, with one cluster appearing to identify as bone marrow-derived macrophages (**BMDM**) while the other expressing markers associated with Trem2 high macrophages. Trem2 high macrophages are characterized by activating complementary pathways, expressing CD9 and extracellular matrix remodeling factors such as cathepsins [43]. CD9 expression inhibits LPS-stimulated macrophage activation resulting in immunosuppression [48]. Cathepsins are proteases that act as extracellular matrix modeling factors, and their expression in macrophages promotes tumor progression and invasion [49, 50]. Trem2 high macrophages may also increase resistance to immune checkpoint blockade therapies [44]. Both macrophage clusters expressed increased M1-like genes and immune activation features during LGG; however, in HGG, immune suppressive factors were significantly overexpressed. In addition, we identified activated microglial populations expressing *Ccl3* and *IL-1b* only in LGG and HGG, but not in NT. Others have described that microglia stimulated with lipopolysaccharide (**LPS**) significantly increased expressing chemokines (e.g., *Ccl3* and *IL-1b*) [51]. Our results suggest that microglia may also be sensitive to changes within the microenvironmental milieu caused by tumorigenesis and non-resident immune cell infiltration resulting in their activation.

However, while differential expression analysis provides understanding of the average fold change of genes between states, gene expression may largely vary within a group or be expressed by a percentage of cells at the single-cell level. Thus, to evaluate whether these effects were representative of a phenotypic shift of an immune cluster during malignant progression or a subset of cells which change during progression, we performed a series of *in-silico* experiments using a random-forest based approach to see if state origin could be accurately predicted and was driven by immunomodulatory signatures at a single-cell level. Surprisingly, all reported immune cell clusters were highly predictive by our model (**Supplementary Figures 6D, 6E, 6F and 10**). Moreover, the immunomodulatory signals identified from our differential expression analysis were key drivers of the model in both macrophage clusters; however, in T-cells, this relationship was less apparent (**Supplementary Figure 5**). Machine learning data for monocytes, CD4-specific T cells, CD8-specific T cells, B cells, NK cells, and immature erythrocytes were additionally not reported due to the low total cell counts of these groups (**Supplementary Figure 2B**). Most interestingly, assaying the top 30 features (genes) which drove stratification of progression state for microglia showed that the same immunomodulatory factors which characterized macrophage cluster 1 and 2 were involved, including CD74 (**Supplementary Figure 10**), implicating a homology in the phenotypic shift of both macrophages and microglia during glioma malignant progression.

In other studies, similar observations of combined signatures for immune activation and suppression factors were observed in microglia and macrophages in the glioma microenvironment [52]. Since both the macrophage clusters expressed increased immunosuppressive factors as malignant progression emerged, we identified additional targets that require further interrogation to determine whether selective depletion or impairment will alter their immunosuppressive macrophage functionality. To that end, we identified *Id2* being upregulated in LGG and HGG for both macrophage clusters. The *Id2-Kdr* axis has been shown to drive myeloid-derived proangiogenic signaling during progression of glioma [13]. Similarly, *Cd74* and *Mif* were both upregulated in LGG and HGG macrophage clusters 1 and 2. GBM patients with MIF-positive tumors had 42% reduced overall survival compared to patients with MIF-negative tumors [53]. In addition, CD74-MIF signaling can block both M1 polarization and IFN-g release in microglia and pharmacological inhibition of CD74 reduced tumor burden in a murine model of high-grade glioma [54].

Furthermore, Ibudilast, which targets MIF signaling, has shown to sensitize glioblastoma cells to temozolomide in a patient-derived glioma xenograft model and may serve as a way to target tumor associated macrophage clusters [53]. Also, monocytic MDSCs express the MIF receptor for CD74 and targeting the MIF-CD74 interaction using Ibudilast ameliorated immunosuppression and amplified CD8+ T cell activity in a GL261 glioma model [55].

Our scRNA sequencing studies also revealed significantly less T cell infiltration in HGG than LGG, making these therapeutic approaches warrant further investigation. In line with this finding, patients with GBM present with extremely low T cell counts, and these T cells are often sequestered in the bone marrow due to the loss of S1P1 in T cells [56]. However, another study reported significant CD8+ T cell infiltration in patients with grade II glioma compared with grade III and IV [57] [58], and higher CD8+ T cell infiltration correlated with a better prognosis in primary GBM [59]. Overall, a significant reduction in T cell counts and increased co-inhibitory molecule expression in the remaining T cells depict a dynamic immunosuppressive and permissive TME within HGGs and during progression.

A limitation of this study is that single-cell transcriptomic analyses focused on isolated CD45+ cells and this selected population comprises only a fraction of immune cells within the TME. However, this isolation process was necessary to capture a sufficient number of immune cells that supported enhanced immune cell cluster resolution at the single cell level. In subsequent studies, we aim to understand the tumor and immune cell cross-talk before and during malignant progression. It will be critical to understand if there are transcriptional commonalities between tumor cells in LGG, intermediate, and HGG time points and their associated myeloid cells, and how this impacts adaptive cell trafficking and activation. In addition, longitudinal studies aimed at immune cell monitoring in peripheral blood from patients may enhance our ability to develop immune-oncologic surveillance strategies that may be predictive of progression and treatment response. Our data suggest that macrophages demonstrate features of immune activation in the low-grade stage and acquire immunosuppressive signatures as the tumor transforms to high-grade. Functional studies targeting macrophages in the dynamic period between low and high-grade glioma may abrogate immunosuppressive mechanisms and provide new therapeutic windows and opportunities to prevent malignant progression.

