## Supplementary for "Single-cell RNA Sequencing Reveals Immunosuppressive Myeloid Cell Diversity and Restricted Cytotoxic Effector Cell Trafficking and Activation During Malignant Progression in Glioma"

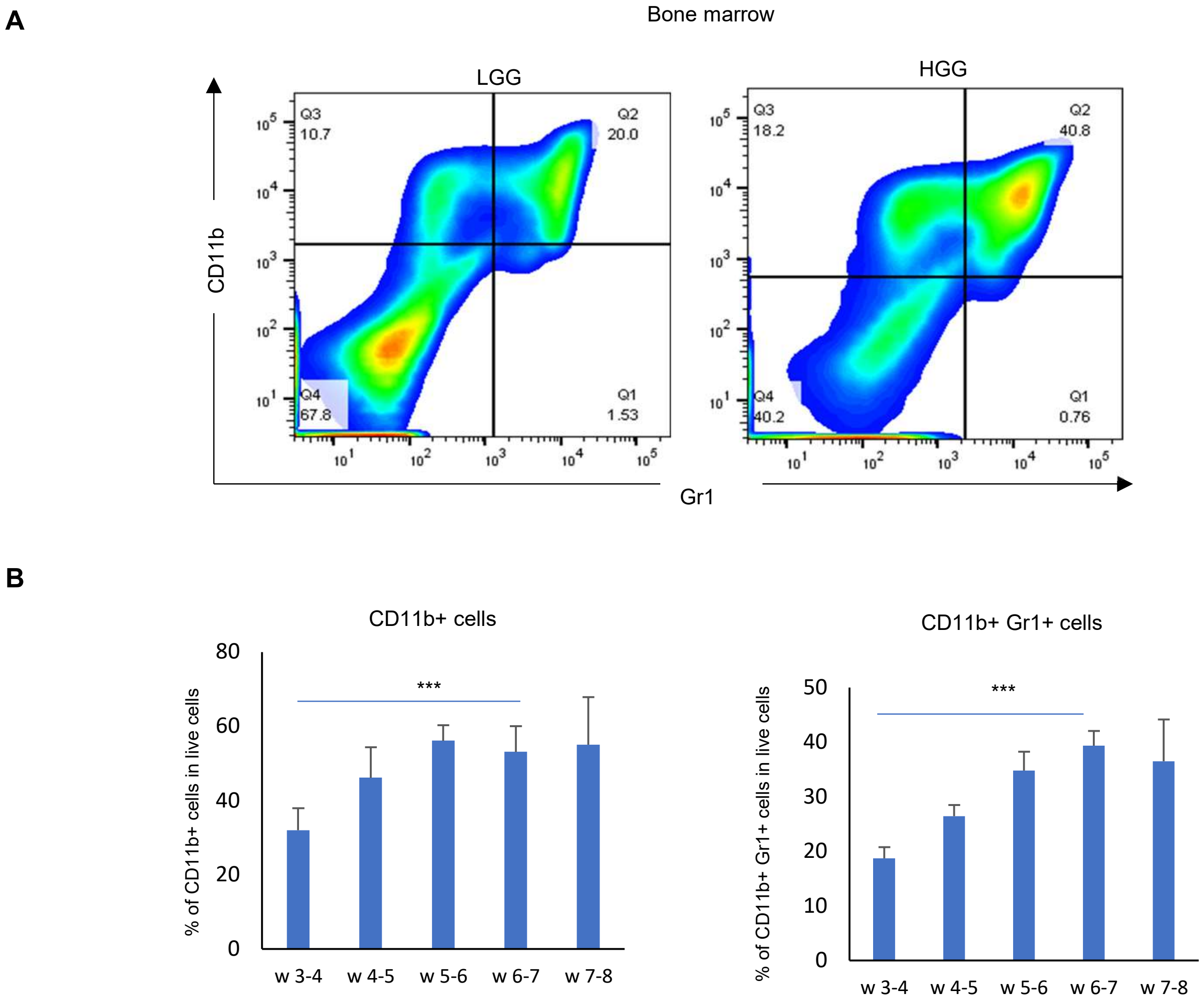


**Supplementary Figure 1. Increased myeloid-derived cells in the bone marrow of animals bearing high-grade glioma (HGG).** (**A**) Representative flow cytometry staining for CD11b and Gr1 in the bone marrow of RCAS animals at specified time points, w 3-4 (n=3), w 4-5 (n=5), w 5-6 (n=3) and w 6-7 (n=6) (**B**) Quantification of percentage of CD11b+ cells and CD11b+Gr1+ cells in bone marrow. Two-tailed unpaired student's t-test was used to calculate statistical significance. ***P<0.005

### Supplementary Figure 2

A

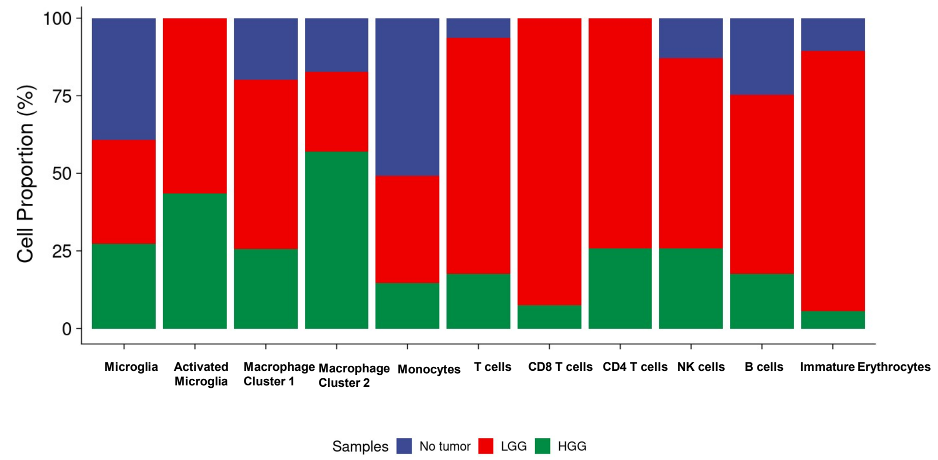


B

| Cluster Nameaaaaaaaa | No Tumor | LGG | HGG | Total |
|---|---|---|---|---|
| Microglia | 2713 | 2314 | 1879 | 6906 |
| Activated Microglia | 0 | 453 | 348 | 801 |
| Macrophage Cluster 1 | 302 | 829 | 389 | 1520 |
| Macrophage Cluster 2 | 397 | 588 | 1306 | 2291 |
| Monocytes | 28 | 19 | 8 | 55 |
| T cells | 33 | 390 | 90 | 513 |
| CD4 T cells | 0 | 23 | 8 | 31 |
| CD8 T cells | 0 | 147 | 12 | 159 |
| B cells | 49 | 114 | 35 | 198 |
| NK cells | 9 | 43 | 18 | 70 |
| Immature Erythrocytes | 13 | 104 | 7 | 124 |
| Total | 3544 | 5024 | 4100 | 12668 |

C

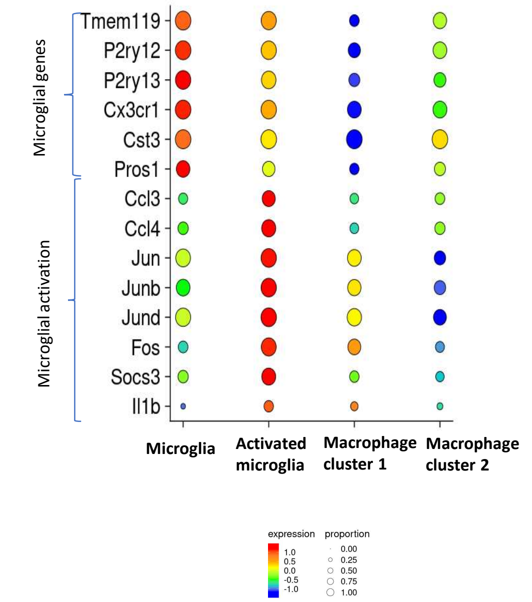


D

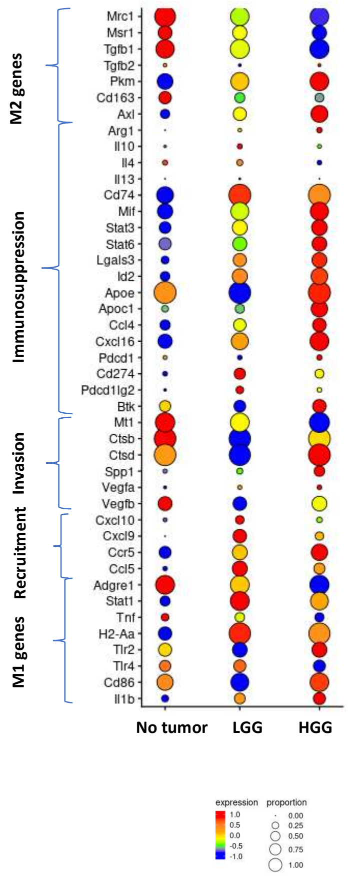


E

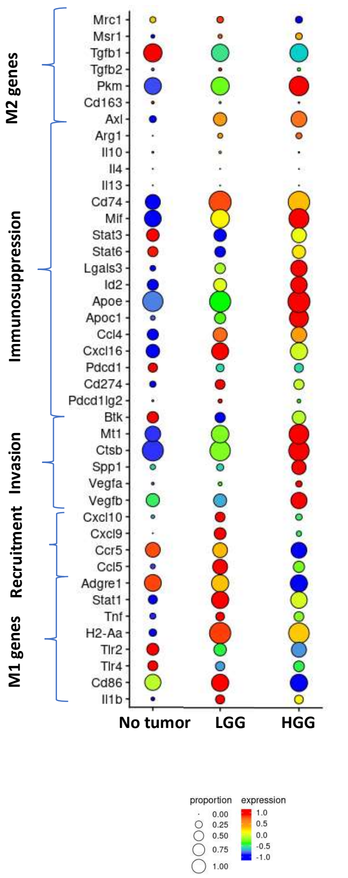


**Supplementary Figure 2. Low grade glioma sample constitute the highest proportion of cells in activated microglia, macrophage cluster 1, T cells, CD4 T cells, CD8 T cells, NK and B cells** (A) Percentages of cell counts of each cell cluster differentiated by sample type, no tumor (blue), low grade glioma (red), high grade glioma (blue). (B) Table with absolute cell counts for each cluster observed at different tumor time points. (C) Differential expression analysis for canonical microglial genes (Tmem119, P2ry12, P2ry13, Cx3cr1, Cst3, Pros1), genes associated with microglial transcriptional activation (Jun, Junb, Jund, Fos, Socs3) and chemokine secretion (Ccl3, Ccl4 and Il1b) in microglial and macrophage clusters in integrated dataset. Differential expression analysis of genes related to M2 polarization, immunosuppression, invasion, recruitment and M1 polarization in macrophage cluster 1 (D) and macrophage cluster 2 (E). Size of the circle represents proportions of cells expressing the gene and color represents expression level.

Supplementary Figure 3

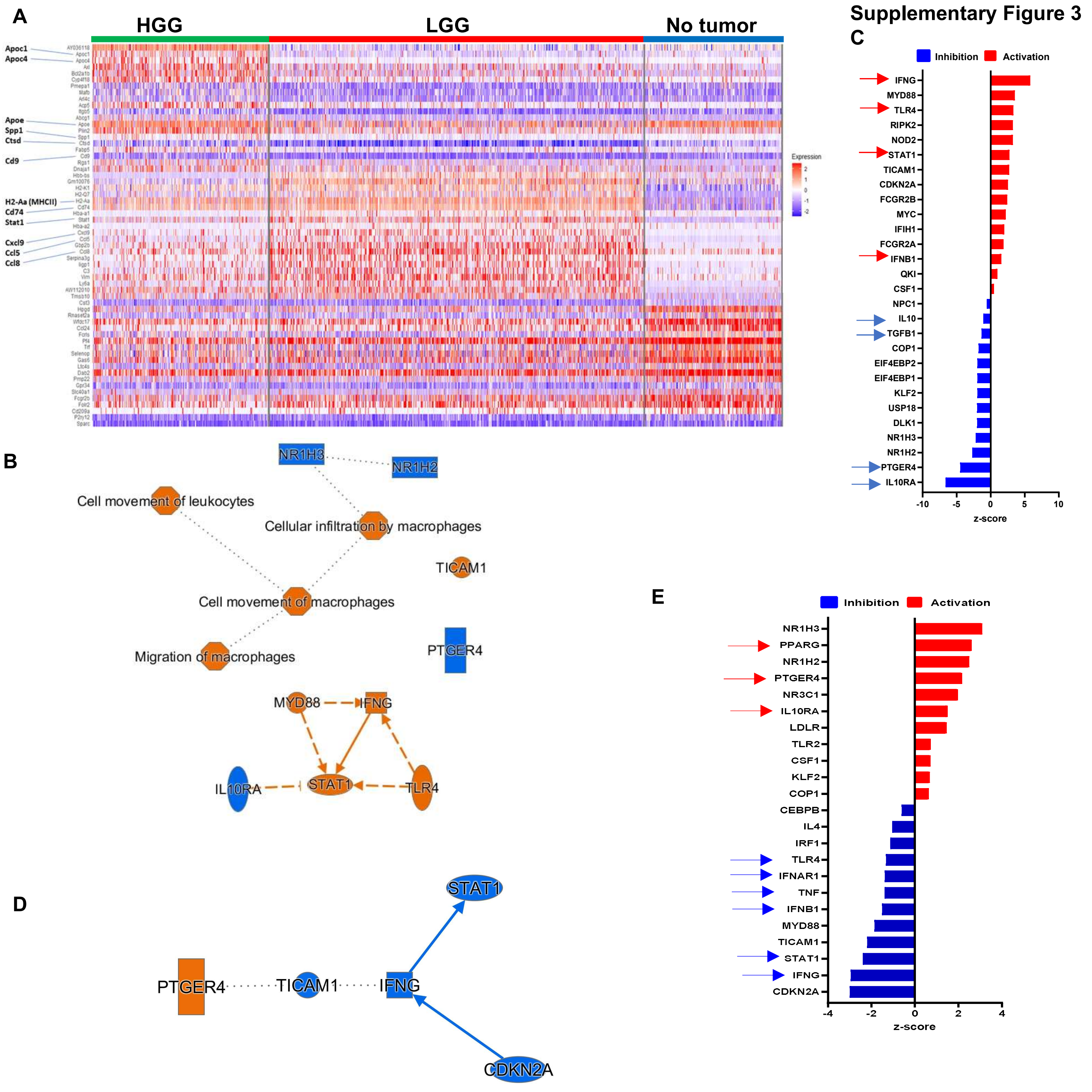


**Supplementary Figure 3. Macrophage cluster 1 in LGG express increased levels of MHCII genes and chemokines associated with T cell recruitment.** (A) Heatmap of top 20 differentially expressed genes in high grade glioma, low grade glioma and no tumor. Highlighted are genes involved in macrophage lipid metabolism (Apoc1, Apoc4, Apoe), immunosuppression (Spp1, Cd74) matrix remodeling (Cstd, Cd9), macrophage activation (MHCII, Stat1) and chemokine secretion (Cxcl9, Ccl5 and Ccl8). Ingenuity pathway analysis (IPA) was performed for macrophage cluster 1 in LGG vs no tumor. (B) represents the networks of pathways activated (orange) and inhibited (blue) in LGG vs NT (C) Top upstream predicted targets with corresponding z-scores, targets upregulated in red and downregulated in blue in LGG vs NT. Ingenuity pathway analysis (IPA) was performed for macrophage clusters for HGG vs LGG. (D) represents the networks of pathways activated (orange) and inhibited (blue) in macrophage cluster 1 in HGG vs LGG. (E) Top upstream predicted targets in macrophage cluster 1 with corresponding z-scores, targets upregulated in red and downregulated in blue in HGG vs LGG.

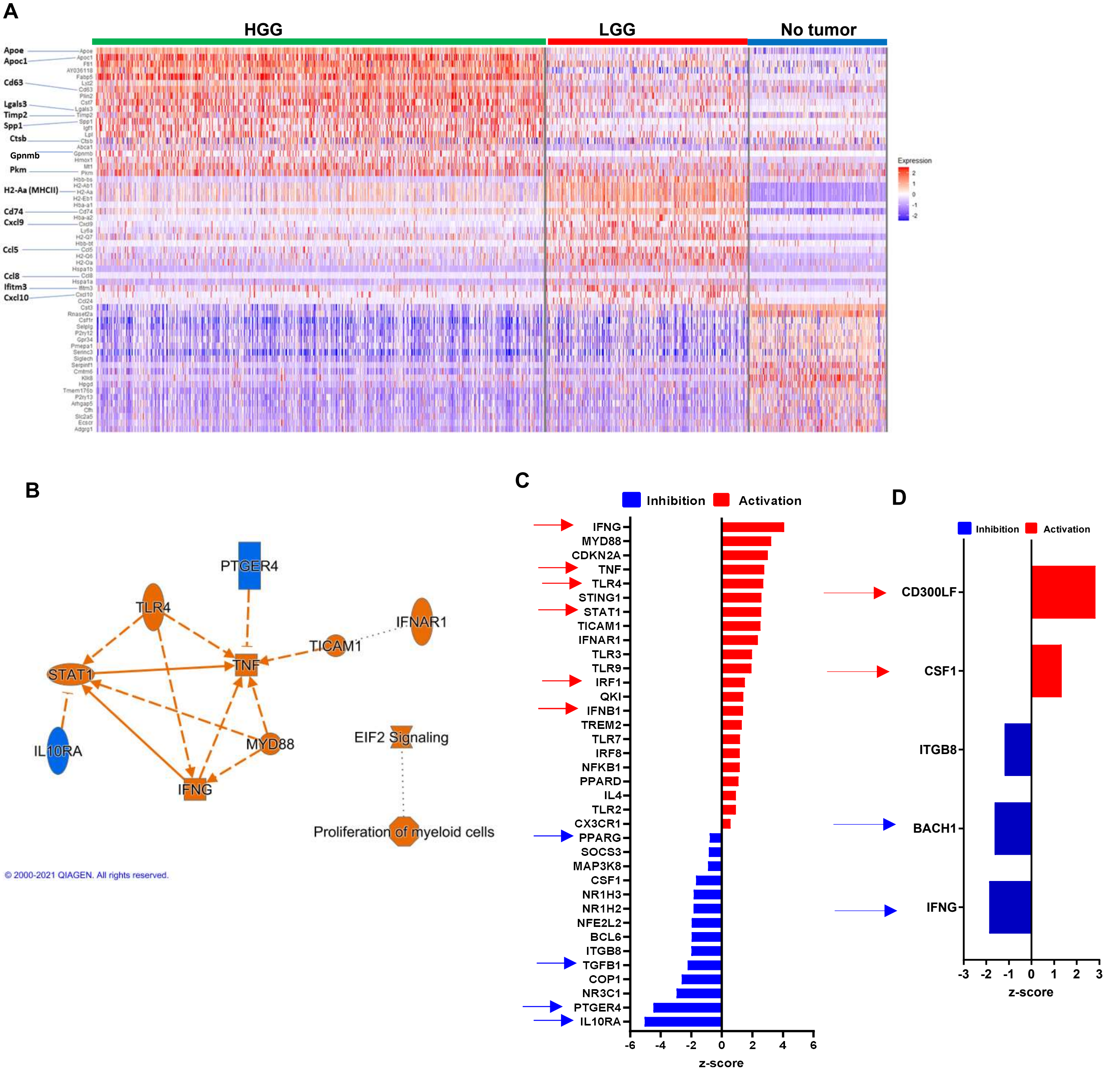


**Supplementary Figure 4. Macrophage cluster 2 in LGG express increased levels of MHCII genes and chemokines associated with T cell recruitment.** (A) Heatmap of top 20 differentially expressed genes in macrophage cluster 2 between high grade glioma, low grade glioma and no tumor. Highlighted are genes associated with macrophage lipid metabolism (Apoe, Spp1), Trem2 signature (Cd63, Cd9, Timp2, Ctsb), immunosuppression (Spp1, Pkm and Cd74) macrophage activation (MHCII, Stat1) and chemokine secretion (Cxcl9, Ccl5, Ccl8 and Cxcl10). Ingenuity pathway analysis (IPA) was performed for macrophage cluster 2 in LGG vs no tumor. (B) represents the networks of pathways activated (orange) and inhibited (blue) in macrophage cluster 2 in LGG vs NT. Top upstream predicted targets with corresponding z-scores, targets upregulated in red and downregulated in blue in LGG vs NT (C) and HGG vs LGG (D).

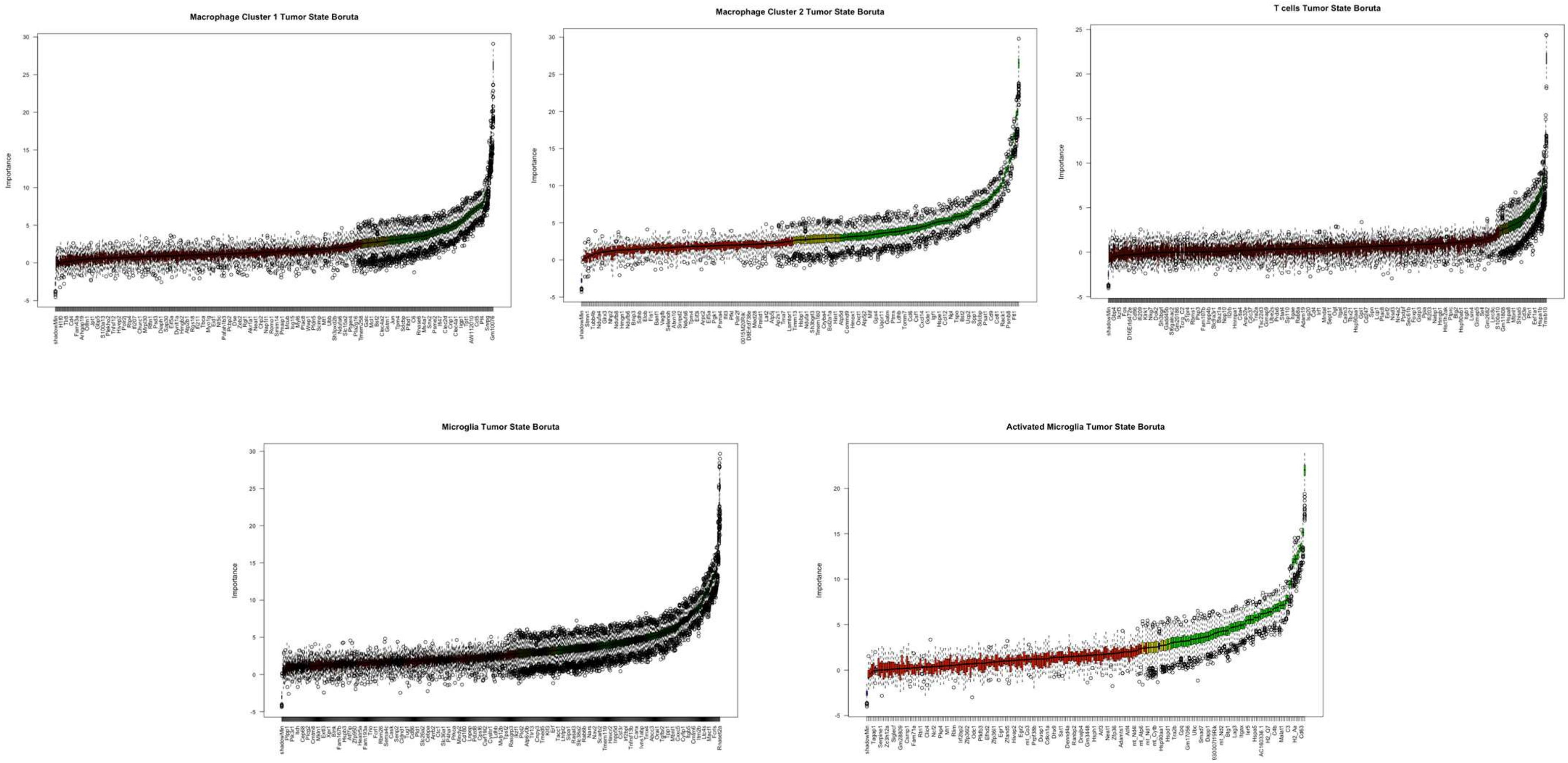


**Supplementary Figure 5. Boruta feature reduction removes several non-predictive genes in all cell cluster types.** Prior to random forest machine learning classification, features (genes) are reduced to eliminate features which are likely non-predictive of discerning origin of cancer progression for a cell by comparing features to randomized shadow features (blue). Accepted features which significantly outperform shadow features are accepted (green) and those that fail are rejected (red). Yellow features are indeterminate after 500 permutations used to evaluate importance. Exported features for machine learning modeling are all non-rejected features (green + yellow).

**Supplementary Figure 6**

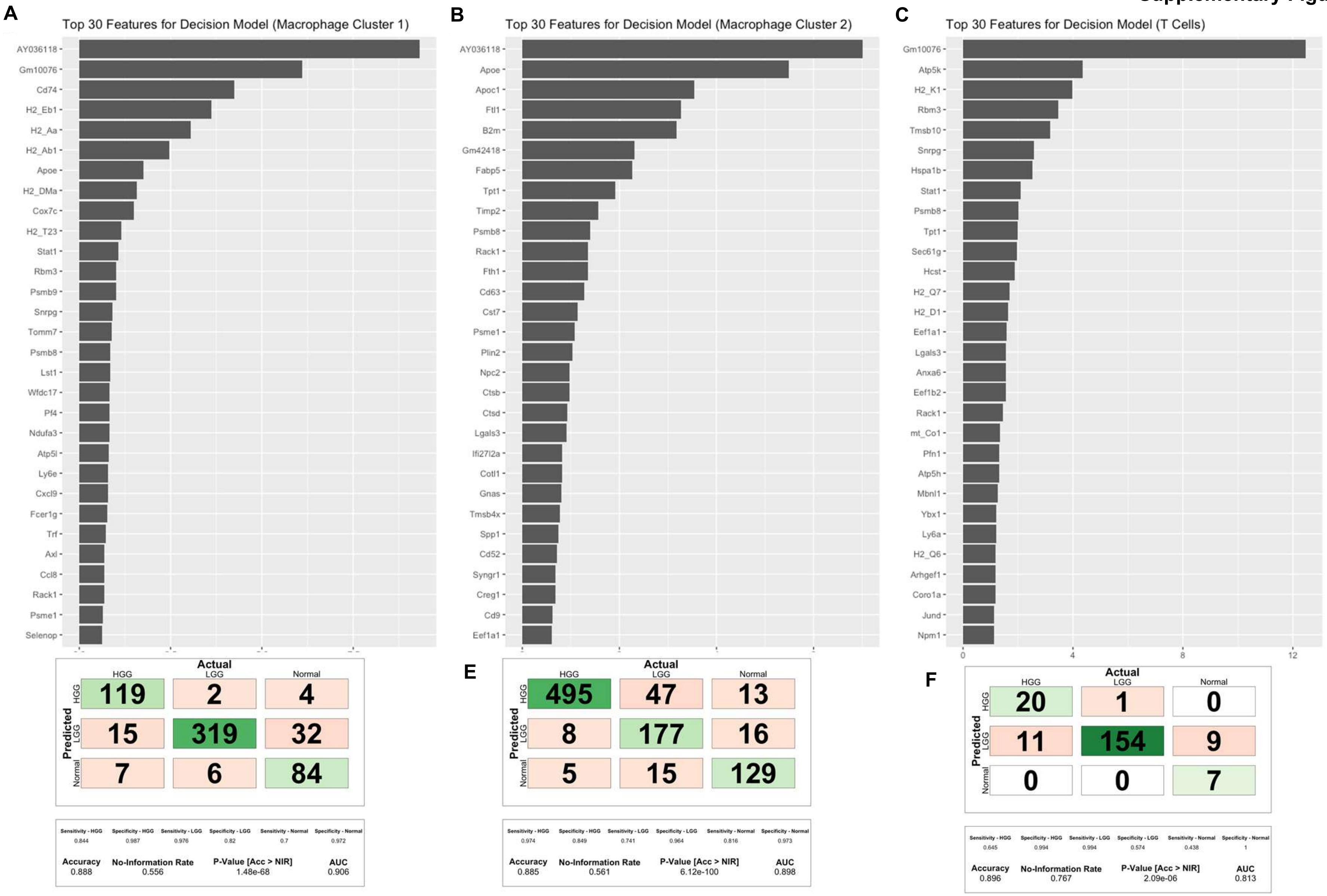


**Supplementary Figure 6. Random forest models of macrophage cluster 1 & 2 are driven by immunomodulatory markers, but T cells are not.** Top 30 ranked features were extracted from machine learning models built to predict cancer progression state origin for cell from (A) Macrophage Cluster 1, (B) Macrophage Cluster 2, and (C) T Cells. Notably, genes identified as differentially expressed with immunomodulatory functions were seen as key drivers of model performance for macrophage cluster 1 (e.g. CD74, MHC genes, ApoE) and macrophage cluster 2 (e.g. ApoE, ApoC1). However, features (genes) driving the T cell model contained did not show many immunomodulatory drivers likely due to heterogenous cell subtypes. Confusion matrix scores of the validation test-set from cells in macrophage cluster 1 (D), macrophage cluster 2 (E) and T cells (F).

**Supplementary Figure 7**

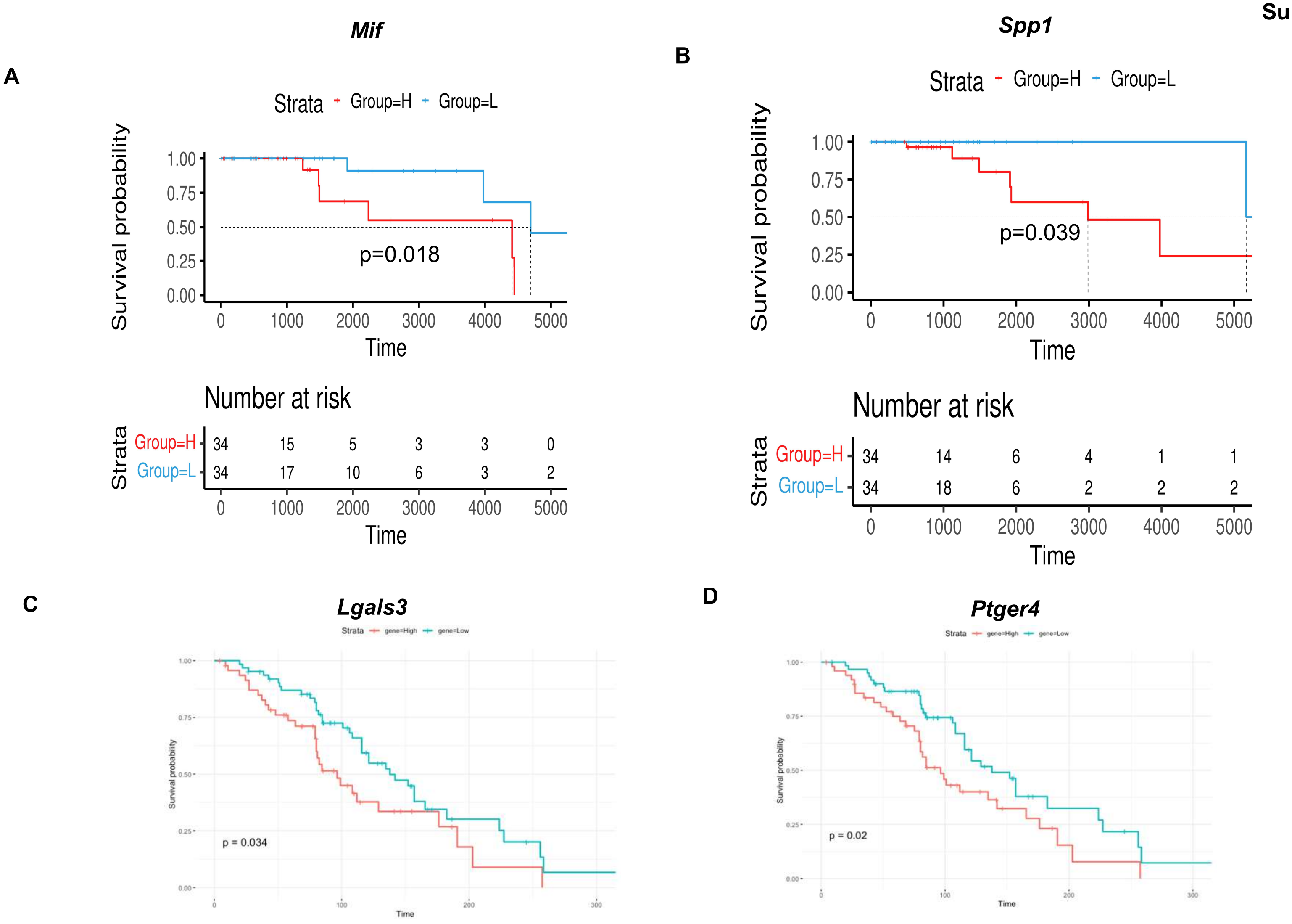


| | n | events | median | 0.95LCL | 0.95UCL |
|---|---|---|---|---|---|
| gene=High | 48 | 29 | 96.4 | 80.1 | NA |
| gene=Low | 63 | 37 | 137.8 | 115.7 | 224 |

High z score >0, Low z score <0

| | n | events | median | 0.95LCL | 0.95UCL |
|---|---|---|---|---|---|
| gene=High | 50 | 33 | 96.4 | 80.1 | 165 |
| gene=Low | 61 | 33 | 137.8 | 115.7 | 227 |

High z score >0, Low z score <0

**Supplementary Figure 7. Reduced overall survival in glioma patients expressing increased levels of *Mif, Spp1, Lgals3 and Ptger4.*** Overall survival of grade II glioma patients expressing low levels (blue) and high levels (red) of *Mif* and *Spp1* (A and B) and low- and high-grade glioma patients expressing low levels (blue) and high levels (red) of *Lgals3 and Ptger4* (C and D).

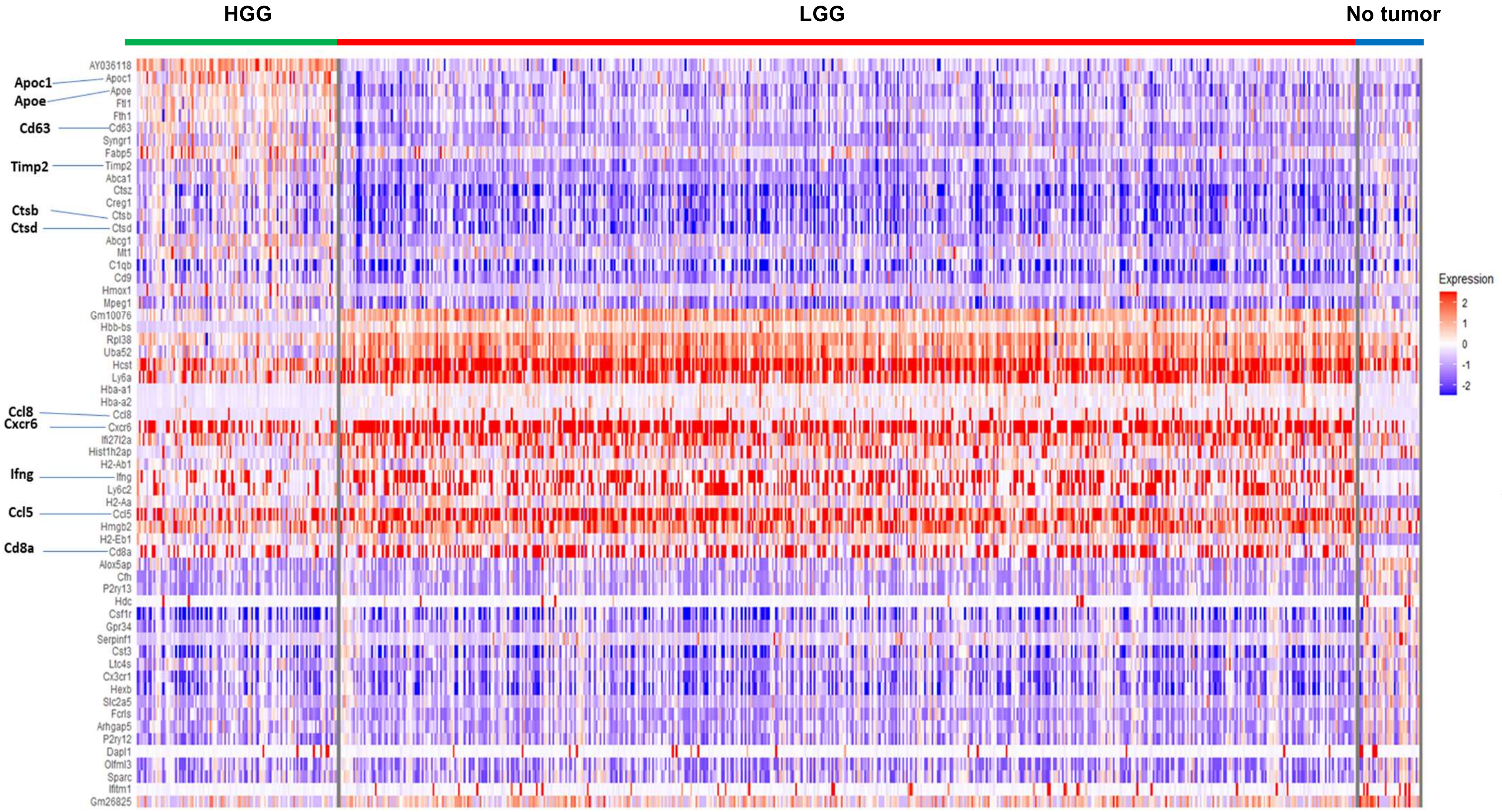


**Supplementary Figure 8. T cells in LGG exhibit increased signatures of immune activation.** Heat map of top 20 differentially expressed genes in CD3 T cells between high grade glioma, low grade glioma and no tumor. Highlighted are genes associated with T cell lipid metabolism and immunosuppression and chemokines involved in recruitment.

**Supplementary Figure 9**

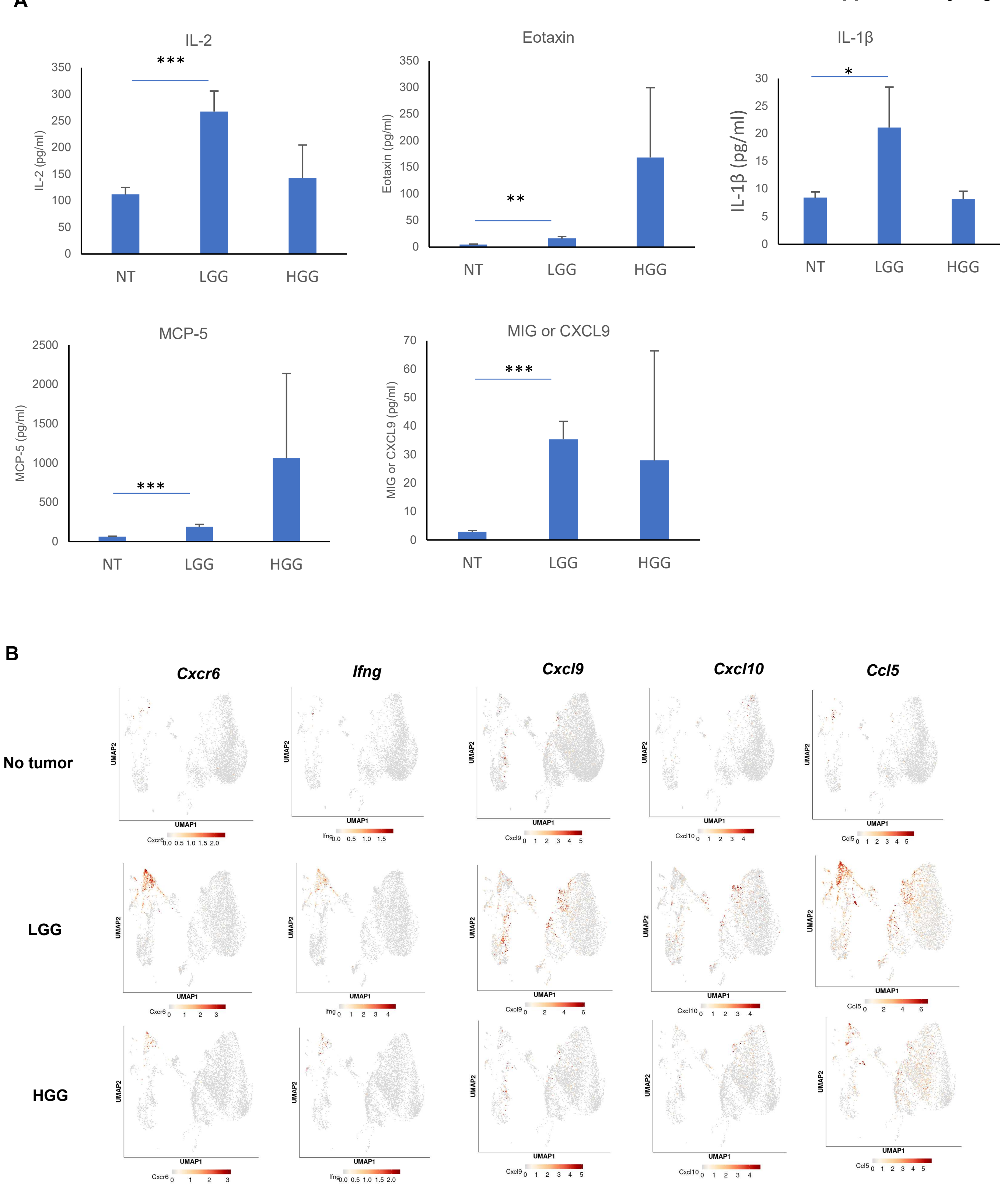


**Supplementary Figure 9. LGG had the highest expression of recruitment factors and Ifng.** (A) Luminex analysis of chemokines IL-2, Eotaxin, IL-1b, MCP-5 and CXCL9 from tissue or tumor lysates from no tumor (n=3), LGG (n=3) and HGG (n=4). (B) Feature plots representing the expression of *Cxcr6, Ifn-g, Cxcl9, Cxcl10* and *Ccl5* across no tumor, LGG and HGG. Statistical significance was calculated using two-tailed unpaired student's t test. * P<0.05, ** P<0.01 and *** P<0.005.

**Supplementary Figure 10**

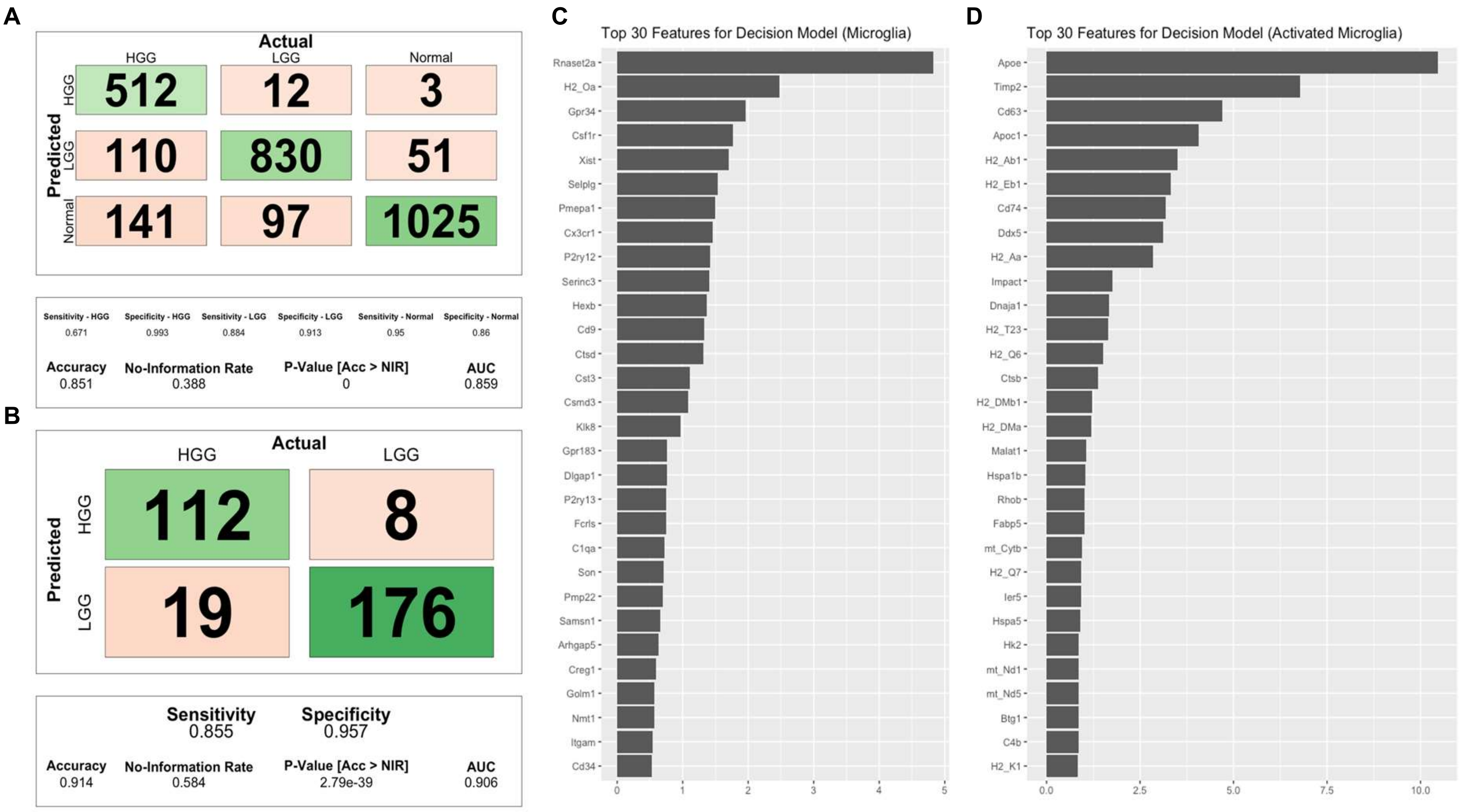


**Supplementary Figure 10. Activated microglia are stratified by immunomodulatory markers shared between macrophage cluster 1 & 2.** Random forest-based machine learning was evaluated for both (A) Microglia and (B) Activated Microglia. Both populations of cells could be stratified with high accuracy between different states of progression**.** However, unlike the primary drivers of model accuracy in (C) Microglia, (D) Activated Microglia showed several markers which had been exhibited to be differentially expressed in macrophages (e.g. ApoE, Timp2, CD63, ApoC1, CD74).

| Characteristic | Total (N=236)* | IDH Mutation and 1p/19q Codeletion (N=64) | IDH Mutation and No 1p/19q Codeletion (N=152) | IDH Wild Type (N=19) |
|---|---|---|---|---|
| **Histologic Type** | | | | |
| **Oligodendroglioma** | | | | |
| **Grade II** | 49 | 29 | 17 | 3 |
| **Grade III** | 21 | 12 | 6 | 3 |
| **Astrocytoma** | | | | |
| **Grade II** | 44 | 1 | 36 | 7 |
| **Grade III** | 55 | 2 | 48 | 4 |
| **Mixed Glioma** | | | | |
| **Grade II** | 42 | 13 | 29 | 0 |
| **Grade III** | 25 | 7 | 16 | 2 |
| **Age at Diagnosis** | | | | |
| **Mean** | 31.31 | 32.70 | 31.07 | 28.68 |
| **Range** | 14-39 | 17-39 | 14-39 | 21-38 |
| **Gender** | | | | |
| **Female** | 102 | 26 | 64 | 12 |
| **Male** | 134 | 38 | 88 | 7 |
| **Race** | | | | |
| **White** | 219 | 56 | 145 | 17 |
| **Black/African American** | 7 | 3 | 4 | 0 |
| **Asian** | 3 | 2 | 1 | 0 |
| **Not Reported** | 7 | 3 | 2 | 2 |
| **Ethnicity** | | | | |
| **Hispanic** | 19 | 7 | 11 | 1 |
| **Non-Hispanic** | 200 | 53 | 131 | 15 |
| **Not Reported** | 17 | 4 | 10 | 3 |
| **Mutational Status** | | | | |
| **MGMT** | | | | |
| **Methylated** | 197 | 63 | 131 | 2 |
| **Unmethylated** | 39 | 1 | 21 | 17 |
| **TERT** | | | | |
| **Mutant** | 36 | 33 | 2 | 1 |
| **Wild Type** | 105 | 1 | 90 | 14 |
| **NA** | 95 | 30 | 60 | 4 |
| **ATRX** | | | | |
| **Mutant** | 117 | 1 | 112 | 4 |
| **Wild Type** | 118 | 63 | 40 | 15 |
| **NA** | 1 | 0 | 0 | 0 |
| **BRAF V600E** | | | | |
| **Mutant** | 1 | 0 | 0 | 1 |
| **Wild Type** | 234 | 64 | 152 | 18 |
| **NA** | 1 | 0 | 0 | 0 |

***Note: 1 Sample did not have IDH status. Known profile was Grade III Astrocytoma in white, non-Hispanic male, age 32. Patient deceased after 1183 days. Mutational profile was 1p/19q co-deleted, MGMT methylated. Missing factors were all not reported.**

**Supplementary Table 1.** Demographic data and clinical characteristics of patients from TCGA-LGG (under 40 years of age) used for survival analysis.
